# miR-181a Regulates p62/SQSTM1, Parkin and Protein DJ-1 Promoting Mitochondrial Dynamics in Skeletal Muscle Ageing

**DOI:** 10.1101/805176

**Authors:** Katarzyna Goljanek-Whysall, Ana Soriano-Arroquia, Rachel McCormick, Caroline Chinda, Brian McDonagh

## Abstract

One of the key mechanisms underlying skeletal muscle functional deterioration during ageing is disrupted mitochondrial dynamics. Regulation of mitochondrial dynamics is essential to maintain a healthy mitochondrial population and prevent the accumulation of damaged mitochondria, however the regulatory mechanisms are poorly understood. We demonstrated loss of mitochondrial content and disrupted mitochondrial dynamics in muscle during ageing concomitant with dysregulation of miR-181a target interactions. Using functional approaches and mitoQc assay, we have established that miR-181a is an endogenous regulator of mitochondrial dynamics through concerted regulation of Park2, p62/SQSTM1 and DJ-1 *in vitro*. Downregulation of miR-181a with age was associated with an accumulation of autophagy-related proteins and abnormal mitochondria. Restoring miR-181a levels in old mice prevented accumulation of p62, DJ-1 and PARK2, improved mitochondrial quality and muscle function. These results provide physiological evidence for the potential of microRNA-based interventions for age-related muscle atrophy and of wider significance for diseases with disrupted mitochondrial dynamics.

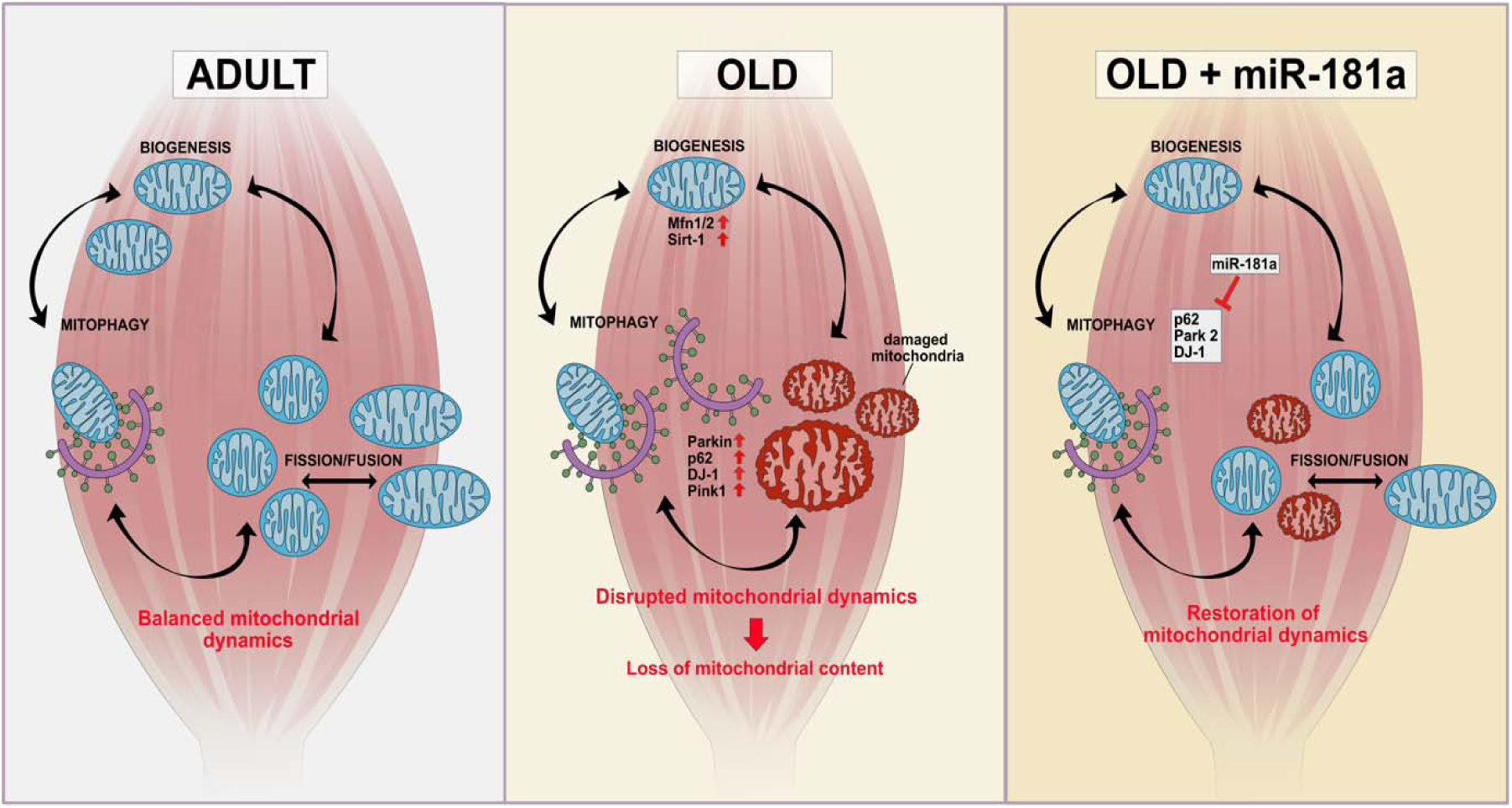

## Introduction

Disrupted mitochondrial dynamics is one of the hallmarks of ageing (Lopez-Otin et al., 2013). Altered mitochondrial morphology and content in skeletal muscle of humans and rodents is one of the pathways consistently associated with age-related loss of muscle mass and function (Bratic and Larsson, 2013; Short et al., 2005), with up to a 30% loss of mitochondrial content reported in fast twitch muscles and an accumulation of dysfunctional mitochondria (Chabi et al., 2008). Moreover, accumulation of damaged or dysfunctional mitochondria increases the oxidation of contractile proteins in muscle; associated with disrupted balance between anabolic and catabolic processes and ultimately sarcopenia (Carnio et al., 2014; O’Leary et al., 2013).

Mitochondria are dynamic organelles that exist in a highly interconnected network and are continually undergoing fusion and fission (for review see (Archer, 2013)). These processes are tightly regulated and necessary to maintain a healthy mitochondrial population by removing and preventing the accumulation of damaged mitochondria. The regulation of mitochondrial remodelling and associated bioenergetic changes, particularly in skeletal muscle, is key for the correct adaptation and response to exercise that results in an increased mitochondrial content with improved fatty acid oxidation and glucose homeostasis (Mansueto et al., 2017). Moreover, the adaptive cellular response of skeletal muscle to exercise requires the autophagic degradation of cellular components, allowing the muscle fibre to rebuild and respond to repetitive bouts of exercise (Vainshtein et al., 2014). Interruption of mitochondrial dynamics, particularly in ageing, can result in mitochondrial swelling, loss of cristae, destruction of the inner membrane and impaired respiration (Bratic and Larsson, 2013). The integration of mitochondrial biogenesis and selective degradation *via* mitophagy is essential for the preservation of healthy muscle, disruption of this balance can result in alterations in muscle bioenergetics and loss of muscle mass and function (Hood et al., 2019). Mitophagy is regulated at numerous levels and a number of distinct mitophagic pathways have been elucidated such as ubiquitin-mediated mitophagy including the Pink/Parkin pathway and ubiquitin independent pathways *via* mitophagy receptors on the outer mitochondrial membrane (e.g. BNIP3), however the exact regulatory mechanisms remain to be fully understood (for reviews see (Montava-Garriga and Ganley, 2019; Palikaras et al., 2018).

microRNAs (miRs) are small 19-25nt long noncoding RNAs that regulate gene expression post-transcriptionally through binding to complementary target sites within mRNAs, usually 3’UTRs, leading to mRNA degradation and/or inhibition of mRNA translation (Bethune et al., 2012). miRs target multiple genes and are considered a robust mechanism of controlling cellular and tissue homeostasis. The role of miRs in the regulation of key cellular mechanisms has become increasingly recognised, including skeletal muscle homeostasis, development, regeneration and atrophy (Cheung et al., 2012; Goljanek-Whysall et al., 2011; Soares et al., 2014). The expression of a number of specific miRs change in skeletal muscle during exercise and ageing (Brown and Goljanek-Whysall, 2015; Hu et al., 2014; Kim et al., 2014; Nielsen et al., 2010). Although limited, functional studies have demonstrated that miRs play a key role in regulating the expression of genes and pathways altered during exercise and/or ageing, contributing to alterations in skeletal muscle mass (Li et al., 2017; Silva et al., 2017; Soares et al., 2014). A number of miRs have been found to be both associated and present within mitochondria, indicating a potential regulatory role (Shen et al., 2016).

In this study, we have demonstrated that age-related disruption of mitochondrial dynamics in skeletal muscle can be improved by restoring the expression of miR-181a-5p (miR-181a). Quantitative proteomics data revealed a reduced mitochondrial protein content with age, concomitant with the upregulation of mitophagy-associated proteins. Ultrastructural analysis of mitochondria revealed abnormal, large mitochondria in muscle during ageing despite increased expression of autophagy, and in particular mitophagy-associated proteins. Parallel analyses of upstream regulators of mitochondrial dynamics identified miR-181a as targeting key autophagy- and mitochondrial dynamics-associated genes*. In vitro* experiments confirmed miR-181a targets p62 and Park2, and demonstrated that miR-181a can regulate mitophagic flux. Restoration of miR-181a content in muscle of old mice *in vivo* prevented accumulation of p62, PARK2 and DJ-1 and preserved mitochondrial content, ultimately resulting in increased myofibre size and muscle force. Together, our data indicate that miR-181a is a potent regulator of muscle mitochondrial dynamics *in vitro* and *in vivo*, providing potential therapeutic avenues for age-related muscle atrophy.

## Results

### Quantitative proteomics reveals decrease in mitochondrial content with age

To characterise changes in the intracellular muscle environment during ageing and associated adaptive response of muscle to contractions, global label-free analysis was used to quantify the overall changes in the proteome of skeletal muscle from quiescent *tibialis anterior* (TA) or TA subjected to 15 min of isometric contractions (mimicking acute exercise) from adult and old mice. Significantly changed proteins (fold change > 2 and −10logP > 20) between quiescent or contracted muscle of adult or old mice demonstrate clear differences in the proteomic content of TA muscle (Fig.1A). Despite detected changes in the abundance of some proteins between contracted muscle of adult and old mice, the major significant proteomic changes detected were as a result of ageing. The most significantly changed pathways in muscle during ageing were: downregulation of mitochondrial proteins and upregulation of contractile apparatus proteins with age (Fig 1B & C). This is consistent with previously reported data showing mitochondrial content decrease in muscle during ageing (Chabi et al., 2008; Hepple, 2014; Smith et al., 2018).

**Figure 1.**
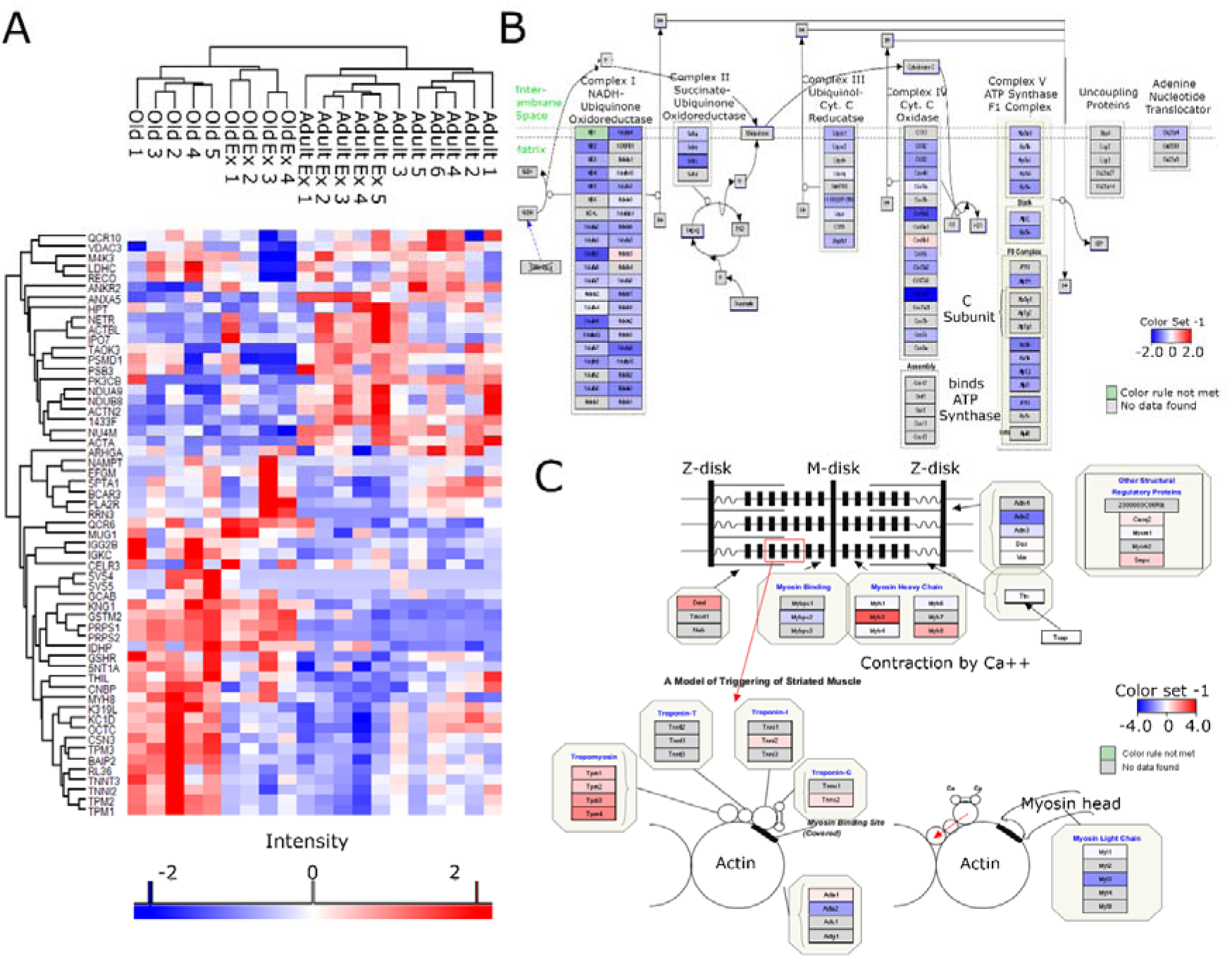
Label free proteomics of TA muscle from adult and old mice. **A.** Heatmap of significantly changed proteins (fold change > 2, −10logP > 20 equivalent to p-value < 0.01) between adult (Adult: 6 months old, n=6), adult exercised (Adult Ex, n=5), old (Old: 24 months old, n=5) and old exercised (Old Ex, n=4). **B, C.** Pathway analysis of quantitative proteomic data reveals a downregulation of proteins involved in mitochondrial electron transport chain and an upregulation of proteins involved in muscle contraction during ageing.

### Autophagy regulatory and effector proteins are more abundant in skeletal muscle from old mice

To further investigate the mechanisms leading to decreased mitochondrial content during ageing, we analysed the expression of known regulators of autophagy and mitochondrial dynamics by western blotting and qPCR. General regulators of the autophagic machinery: Sirtuin 1 (SIRT-1) and Forkhead box protein O3 (FOXO3), as well as effector mitophagy proteins, such as PTEN-induced putative kinase 1 (PINK1), Parkin (PARK2), Protein-DJ-1 (PARK7) and the autophagic adaptor protein p62 (Sequestosome 1, Sqstm1), were upregulated in muscle during ageing (Fig. 2A, B). We also observed downregulation of the expression of Pgc1α with age (Fig. S1). Despite upregulation of autophagy-associated proteins, swollen and abnormal mitochondria were detected by EM in muscle of old mice, suggesting defective mitochondrial dynamics (Fig. 2C). This suggests impaired or dysfunctional mitophagy in skeletal muscle from old compared to adult mice, an increase in p62 levels can be associated with an inhibition of autophagy as it is degraded in cells with normal autophagic flux. (Fig. 2B). Analysis of an autophagy marker LC3 revealed more pronounced LC3 punctae in muscle from old mice (Fig. 2D). The expression of Lc3b was upregulated in the muscle of old mice and the levels of CoxIV and Nd-1, indicators of mitochondrial content, were downregulated in the muscle of old mice, further suggesting dysfunctional autophagic response in ageing muscle and in the adaptation to exercise (Fig. 2E,F). Together, our data demonstrate an accumulation of key regulators of the mitophagic machinery in muscle during ageing and defective mitochondrial dynamics resulting in accumulation of abnormal mitochondria and loss of mitochondrial content. miR-181a, and not miR-181b, miR-181c or miR-181d, was downregulated in TA of mice during ageing and exercise of adult mice only (Fig. 2G, 2H). This suggests that miR-181a is the key miR-181 family member with a role in muscle ageing.

**Figure 2.**
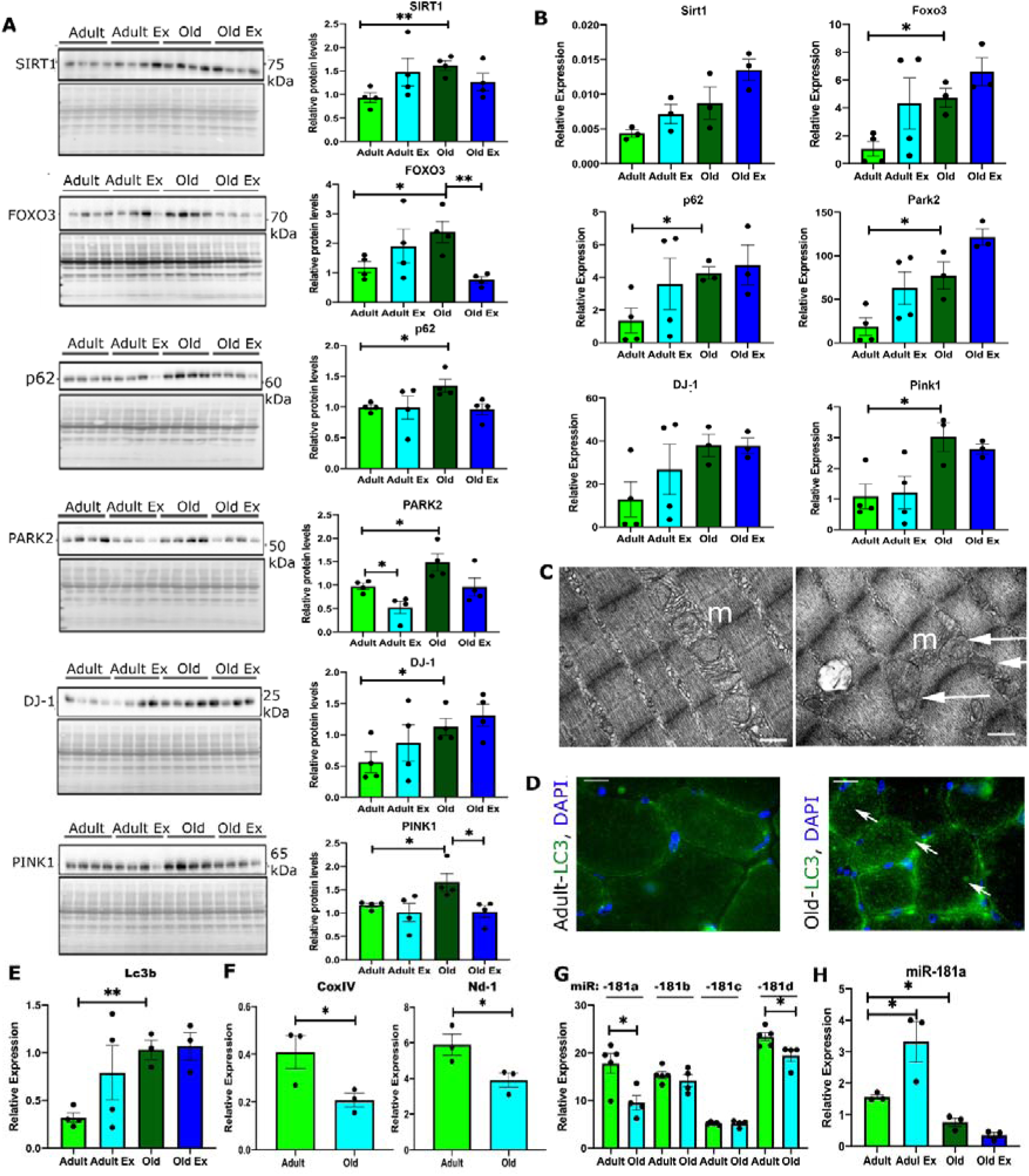
Autophagy and mitochondrial regulators have altered expression in skeletal muscle during ageing. **A.** Western blot analyses showing changes in the protein levels of autophagy-associated proteins in the *tibialis anterior* muscle of adult and old male mice. Levels normalised to Ponceau S (protein) are shown in the quantification graphs (arbitrary units). **B.** Expression levels of mRNA of autophagy-associated gene expression. **C.** Representative EM images of mitochondria from TA muscles of adult and old mice; m – mitochondria; arrows indicate swollen or mitochondria with cristae in disarray. Scale bar: 0.5 µm. **D., E.** Immunostaining (D) and qPCR (E) for LC3 on TA muscle of adult and old mice. Arrows indicate myofibres positive for LC3. Scale bars indicate 100 µm. Arrows indicate LC3 positive fibres. **F.** qPCR analyses of the expression of mitochondrial content markers encoded by the mitochondrial genome. **G.** qPCR analyses of miR-181a, miR-181b, mR-181c and miR-181d expression in the TA of adult and old mice. **H.** qPCR analysis of miR-181a expression in quiescent TA muscle and following contractions protocol of adult and old mice. Adult – 6 months old; old – 24 months old male C57BL6/J mice. Ex – TA following isometric contraction protocol. qPCR: Expression relative to β2-microglobulin (genes) or Rnu6 (miR) is shown. Representative images are shown. n=3-6. Error bars show SEM. * - p<0.05 Student’s t-test.

### miR-181a as putative regulator of mitochondrial dynamics

To determine potential upstream regulators of mitochondrial dynamics we investigated predicted interactions between mitophagy-associated genes and miRs. miR target prediction databases, TargetScan, miRnet and miRWalk, identified miR-181a-5p (miR-181a) as a putative regulator of multiple genes associated with mitochondrial dynamics: previously validated targets (highlighted in bold in Fig. 3A): Park2, Sirt-1, PTEN and Atg-5, and novel putative targets: p62, DJ-1, Mfn1, Mfn2 and Tfam (Fig. 3A). The elevated expression of mitophagy-associated proteins observed in TA from old mice coupled with a decreased expression of miR-181a suggested that miR-181a may act as an important regulator of autophagy and mitochondrial dynamics during muscle ageing.

**Figure 3.**
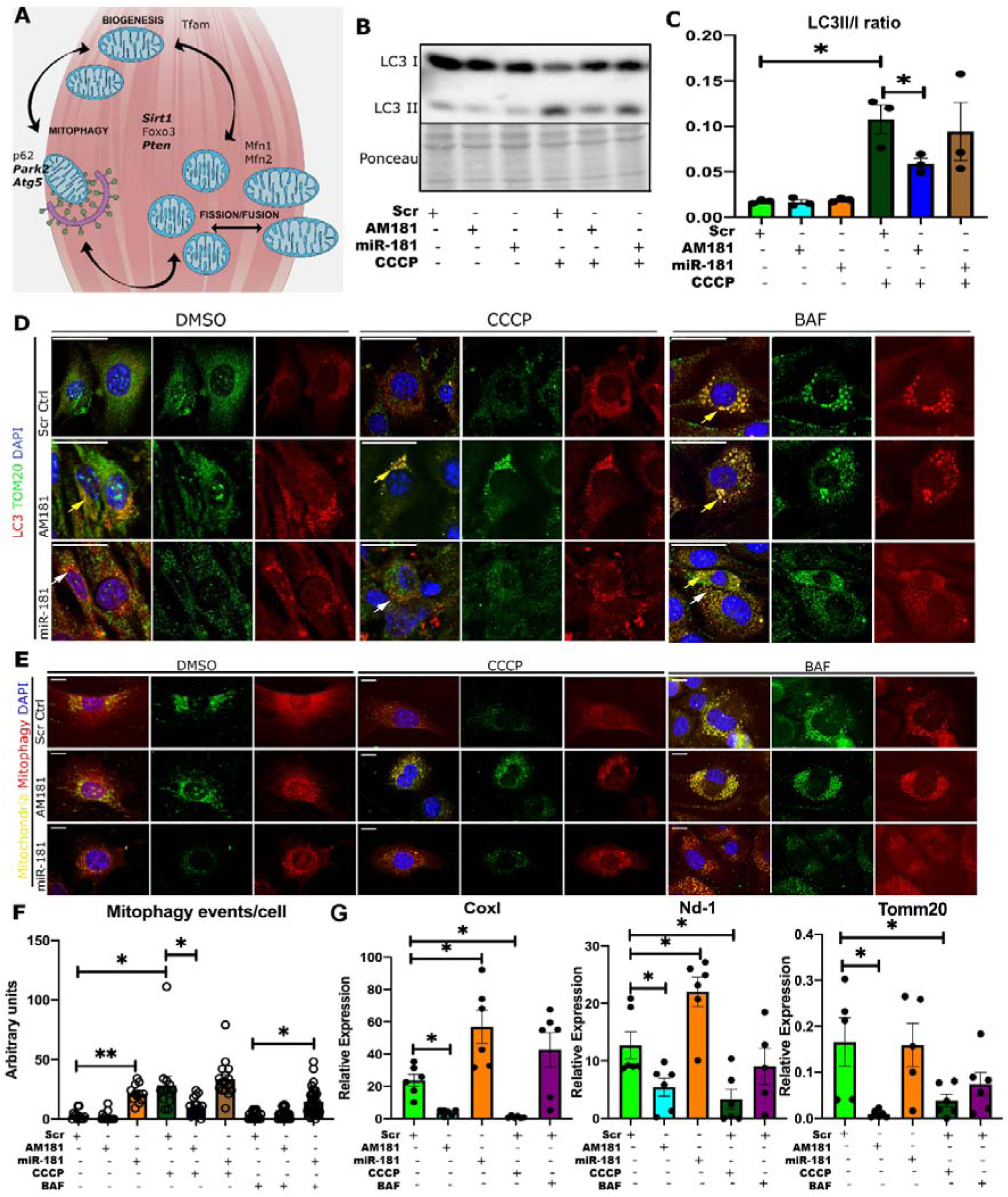
miR-181a regulates mitochondrial dynamics *in vitro*. **A.** Schematic representation of putative and validated (in bold) miR-181a targets associated with mitochondrial dynamics. **B., C.** LC3 I/II western blot and quantification of LC3II/I ratio of C2C12 myoblasts following transfections with scrambled antagomiR, miR-181a mimic or antagomiR-181 and CCCP treatment. **D.** Immunostaining of C2C12 myoblast transfected with scrambled antagomiR (control), miR-181a mimic or antagomiR-181a for LC3 and the mitochondrial marker TOM20 following DMSO, CCCP or BAF treatments. Yellow arrows indicate accumulation of LC3B+TOM20+ punctae and white arrows indicate LC3 positive punctae. Scale bars show 30 µm. Representative images are shown. **E.** Assessment of mitophagic flux in C2C12 cells treated with scrambled antagomiR (control), miR-181a mimic or antagomiR-181a in control (DMSO) or cells treated with CCCP or BAF using mitoQc construct. Mitochondria are labelled with mCherry and GFP; upon mitophagy induction, GFP signal is quenched. Scale bars indicate 10 µm. **F.** Quantification of mitophagic events per cell through analysis of colocalisation of red punctae only. **G.** qPCR analysis of mitochondrial gene expression (Cox I, Nd-1 and Tomm20) in C2C12 myoblasts following DMSO, CCCP and BAF treatments and transfected with scrambled antagomiR (control), miR-181a mimic or antagomiR-181a, respectively, expression relative to β2-microglobulin shown. Error bars show SEM; * - p<0.05 Student’s t-test.

### miR-181a regulates mitochondrial dynamics in C2C12 myoblasts

To investigate whether miR-181a regulates mitochondrial dynamics we used the mitochondrial uncoupler, carbonyl cyanide m-chlorophenyl hydrazone (CCCP) and autophagic flux inhibitor Bafilomycin A1 (BAF) in a C2C12 myoblast model. Treatment of cells with CCCP decreased expression levels of miR-181a, while miR-181a mimic and antagomiR-181a (AM181a) increased and decreased respectively miR-181a levels in C2C12 cells treated with DMSO (control) or CCCP (Fig.S2A). CCCP treatment resulted in formation of LC3 punctae, while BAF treatment resulted in the accumulation of LC3 punctae colocalised with TOM20 (Fig. 3D). miR-181 overexpression resulted in the presence of LC3 positive punctae in DMSO, CCCP and BAF treated cells, whereas inhibition of miR-181a resulted in the accumulation of LC3 and TOM20 colocalised punctae in DMSO, CCCP and BAF-treated cells (Fig. 3D). Moreover, inhibition of miR-181a resulted in decreased LC3 II/ I in myoblasts treated with CCCP (Fig. 3B, C). This suggest inhibition of miR-181a may result in stalled autophagy.

In order to investigate miR-181a-mediated regulation of mitochondrial turnover *via* mitophagy, we used the Mito-QC reporter construct that contains tandem mCherry-GFP tag fused to the mitochondrial targeting sequence of the outer mitochondrial membrane protein, FIS1 (Allen et al., 2013). The mitochondrial network fluoresces red and green under normal conditions but during mitophagy, with the delivery of mitochondria to the acidic environment of lysosomes, GFP fluorescence is quenched while mCherry remains stable (Allen et al., 2013; McWilliams et al., 2016). CCCP treatment increased the number of mitochondria associated with lysosomes in Scr controls which returned to basal levels after BAF treatment (Fig. 3E, F). AM181a samples showed reduced levels of mitophagic events after CCCP treatment (Fig. 3E, F), whereas miR-181a overexpression led to increased mitophagy events in DMSO and BAF-treated cells (Fig. 3E, F). miR-181a or AM181a treatment had no effect on cell viability (cytotoxicity assay) and ATP production was mildly increased in myoblasts treated with miR-181a in the presence of CCCP (Fig. S2B).

To investigate the consequences of mitophagy regulation by miR-181a on mitochondrial content, we analysed the expression of mitochondrial proteins encoded by the mitochondrial genome: Cox I, Nd-1 and encoded by the nuclear genome: Tomm20. The expression of all the mitochondrial genes was increased following miR-181a overexpression and decreased in response to AM181a (Fig. 3G).

### miR-181a regulates the expression of p62, Park2 and DJ-1 in C2C12 myoblasts

We next validated the regulation of miR-181a predicted autophagy-associated targets in an *in vitro* model of mitochondrial uncoupling. No changes in the expression of these proteins were detected following miR-181a overexpression or inhibition in C2C12 myoblasts (Fig. S2 C, D), possibly due to their high turnover. However, in DMSO treated cells, inhibition of miR-181a resulted in increased levels of p62 mRNA and accumulation of p62 protein which did not colocalise with COXIV mitochondrial marker (Fig. 4A, B). miR-181a overexpression resulted in the presence of p62, PARK2 and DJ-1 positive punctae colocalised with the mitochondrial marker COXIV. In CCCP-treated cells, inhibition of miR-181a resulted in accumulation of p62, DJ-1 and Park-2 mRNAs and protein with reduced colocalisation with COXIV. miR-181a overexpression resulted in downregulation of the expression of p62, Park2 and DJ-1 to levels comparable to control cells (Fig. 4A, B). We next analysed changes in the levels of Tfam in myoblasts treated with miR-181a mimic or antagomiR, miR-181a overexpression in C2C12 myoblasts treated with CCCP resulted in upregulation of Tfam expression (Fig. 4B). These results indicate that miR-181a increased mitochondrial turnover *via* mitophagy and potentially concomitant increase in mitochondrial biogenesis as indicated by increase of Tfam expression in C2C12 cells treated with miR-181 and CCCP. This indicates a role for miR-181a in maintaining a population of healthy mitochondria within the cells.

**Figure 4.**
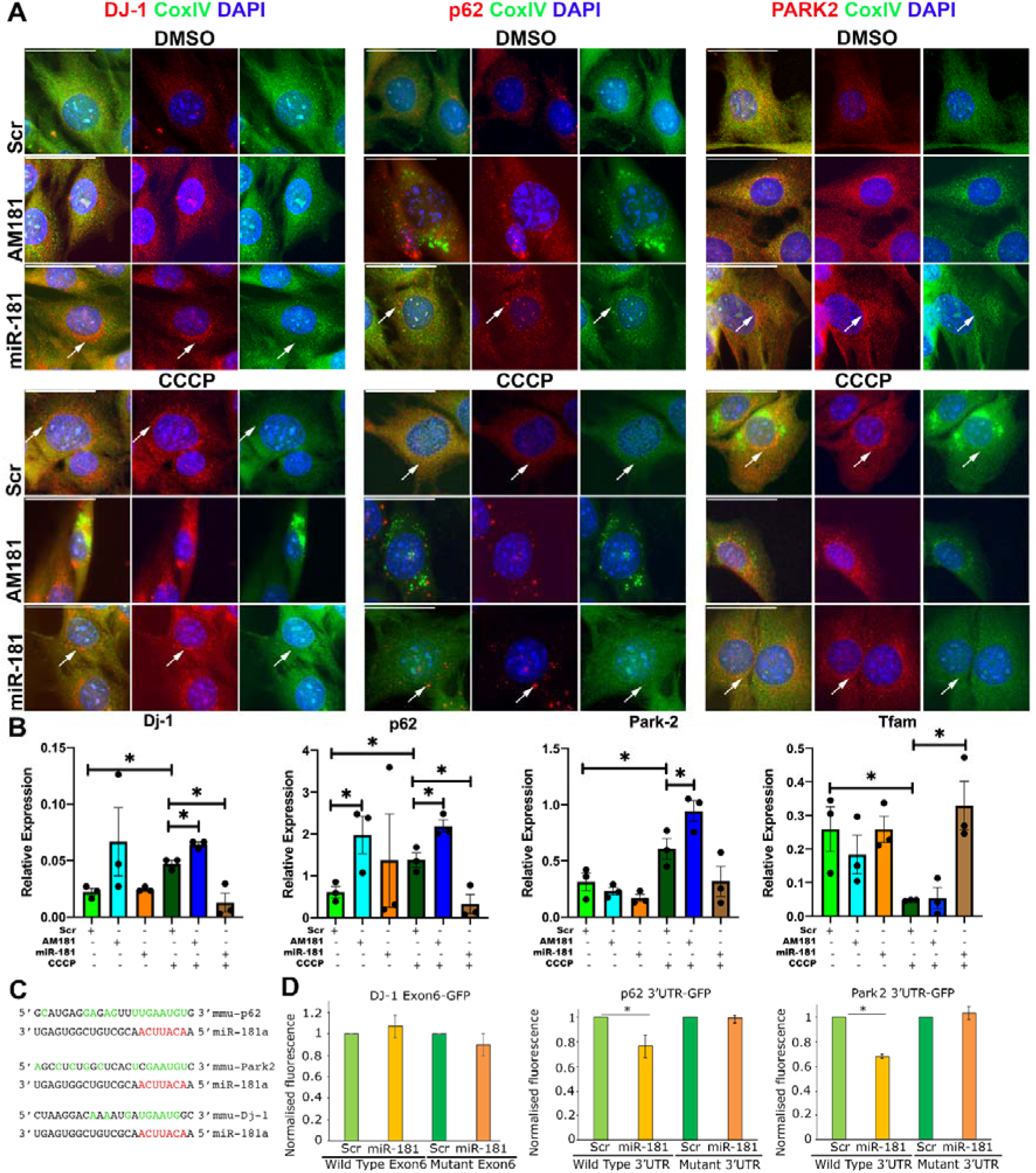
miR-181a targets p62 and Park2 and DJ-1 in C2C12 myoblasts. **A.** Immunostaining for CoxIV (mitochondrial marker) and mitophagy-associated genes: p62, DJ-1 and Park2, showing accumulation of these proteins in C2C12 myoblasts following inhibition of miR-181a function and turnover of mitochondria following overexpression of miR-181a. Arrows indicate regions with protein expression and reduced CoxIV expression indicating mitophagy. Representative images are shown. Scale bars indicate 50 µm. **B.** qPCR showing the effects of miR-181 in C2C12 myoblasts following mitophagy induction on the expression of mitophagy-associated genes, relative to β2-microglobulin. n=3-4. Error bars show SEM. * - p<0.05; unpaired Student’s t-test. **C.** miR-181a predicted binding sites within the 3’UTRs of murine p62 and Park2 and within an exon of murine DJ-1. Red – microRNA seed sequence; green – nucleotides complementary to the microRNAs sequence. **D.** Normalised GFP fluorescence in myoblasts following co-transfection with GFP sensor construct containing wild type or mutated miR-181a predicted binding site and miR-181a mimic or scrambled miRNA mimic. n=3. Error bars show stdev. * - p<0.05 Student’s t-test

We also analysed changes in localisation of BNIP3 and LAMP1 with TOM20 in C2C12 cells treated with miR-181a or AM181a in control, CCCP or BAF-treated cells. Both experiments demonstrated that AM181a treatment leads to accumulation of BNIP3 or LAMP1 punctae colocalised with TOM20, further confirming that low levels of miR-181a are associated with inhibition of mitophagic flux (Fig. S3). However, no miR-181a binding site could be found in the mRNA sequences of Bnip3 or Lamp1.

To investigate direct regulation of autophagy-associated targets by miR-181a, we used 3’UTR of p62 and Park2, DJ-1 Exon 6 in GFP sensor constructs. Although miR binding sites within exons are very uncommon, we analysed this specific putative binding site due to strong effects of miR-181a on DJ-1 expression (Fig. 4C). GFP reporter constructs were transfected into C2C12 cells in the presence of scrambled or miR-181a mimic. miR-181a treatment led to significant decrease in GFP fluorescence from GFP-p62 and GFP-Park2, but not GFP-DJ-1 constructs, as compared to scrambled-treated controls (Fig. 4D). The mutations of miR-181a binding sites rendered all constructs non-responsive to miR-181a overexpression as compared to scrambled controls. This confirms p62 and Park2 as direct, targets of miR-181a, whereas DJ-1 expression may be regulated by miR-181a in an indirect manner (Fig. 4C, D).

### miR-181a regulates expression of p62, DJ-1 and Parkin *in vivo*

The expression and localisation of miR-181a targets were analysed in the muscle of adult and old mice treated with saline, miR-181a mimic or AM181a (Figs. 5, 6 and S6). The muscle of control old mice was characterised by the presence of increased p62, DJ-1 and PARK2-positive myofibres as compared to the muscle of control adult mice confirming disrupted or inhibited autophagy during ageing (Fig. 5A). AM181a treatment of adult mice led to increased number of p62, DJ-1- and PARK2-positive myofibres, whereas fewer p62, DJ-1 and PARK2 positive myofibres were detected in miR-181a-treated old mice compared to their respective controls (Fig. 5A). Inhibition of miR-181a in muscle of adult mice led to accumulation of p62 and DJ-1 mRNA, whereas overexpression of miR-181a in muscle of old mice led to reduced levels of p62, Park2 and DJ-1 mRNA as compared to muscle of control old mice to levels observed in muscle of adult mice (Fig. 5B). This suggests that lower levels of miR-181a in muscle are associated with accumulation of its target genes and dysfunctional mitophagy, potentially leading to accumulation of dysfunctional mitochondria (Fig. 6G). To further investigate the role of miR-181a in regulating mitochondria we analysed the expression of mitochondrial genes (CoxI, and Nd-1) in TA muscle of adult and old mice. Consistently, in muscle of old mice, miR-181a treatment led to increased expression of the mitochondrial genes (Fig. 5C). Furthermore, we investigated changes in expression of transcription factors promoting mitochondrial biogenesis: the master regulator of mitochondrial biogenesis (PGC1α) and the mitochondrial transcription factor (Tfam). Consistently in miR-181a-treated muscle, the expression of these mitochondrial associated genes was upregulated (Fig. 5C, S5). We also observed changes in the expression of Mfn1 in mice treated with AM181a or miR-181a (Fig. S5). Together these results indicate that miR-181a promotes mitochondrial dynamics through concomitant activation of mitophagy and mitochondrial biogenesis in skeletal muscle from adult and old mice.

**Figure 5.**
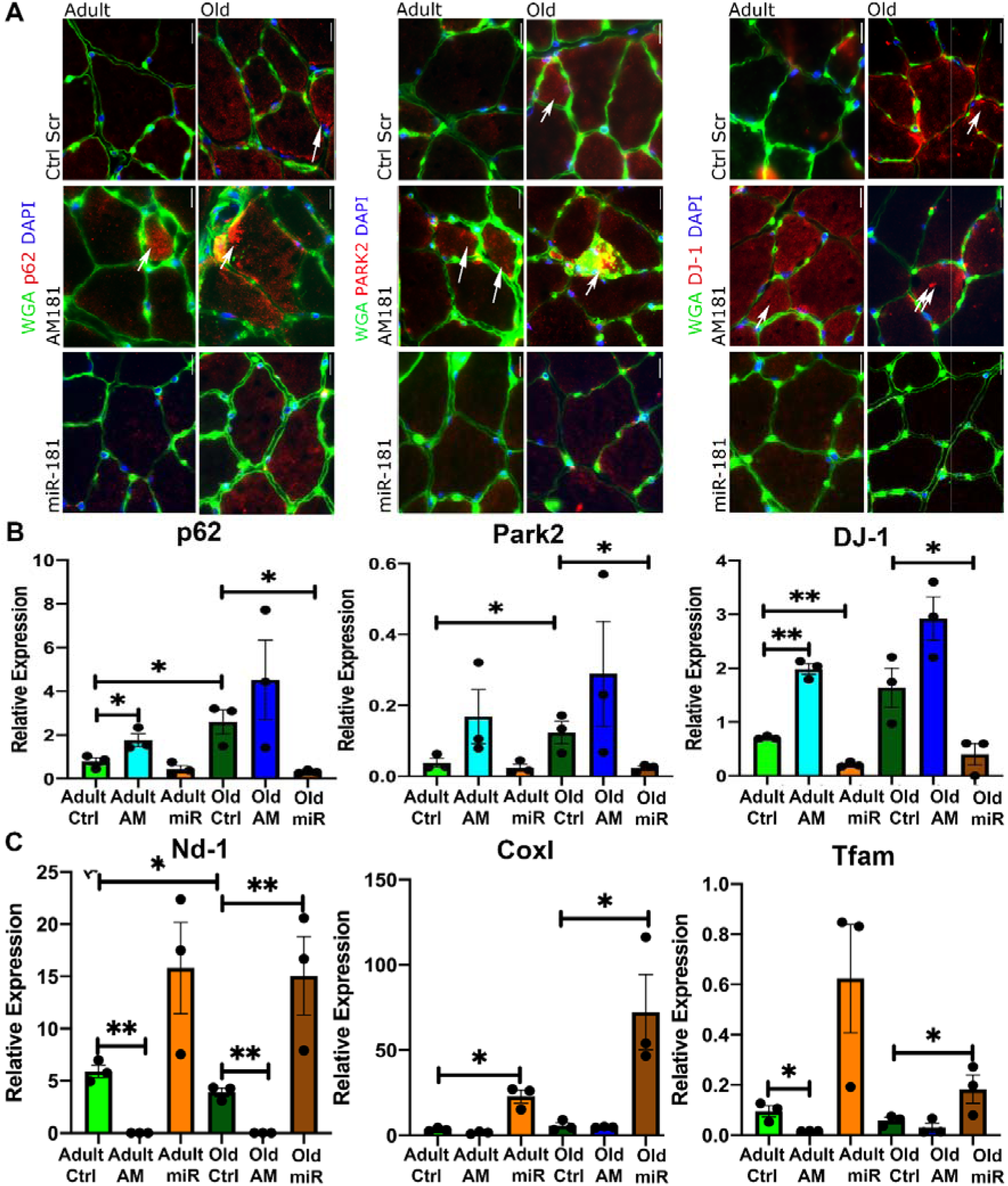
miR-181a regulates expression of p62, DJ-1 and Parkin and affects mitochondrial content markers *in vivo*. **A.** Representative images of TAs immunostained for p62, DJ-1 and Park2 following miR-181a gain- and loss-of-function. Arrows indicate positive fibres. Scale bars indicate 100 µm. n=3. **B.** miR-181a gain- and loss-of-function in TA of adult and old mice leads to changes in the expression of p62, DJ-1 and Park2 mRNA, relative to β2-microglobulin. **C.** miR-181a overexpression increases the expression of mRNA of mitochondrial genes (Cox I, Nd-1) and regulator of mitochondrial biogenesis (Tfam) in TA of adult and old mice, relative to β2-microglobulin. Error bars show SEM * - p<0.05 Student T-test. Adult – 6 months old; old –24 months old male C57BL6/J mice; Ctrl - saline.

**Figure 6.**
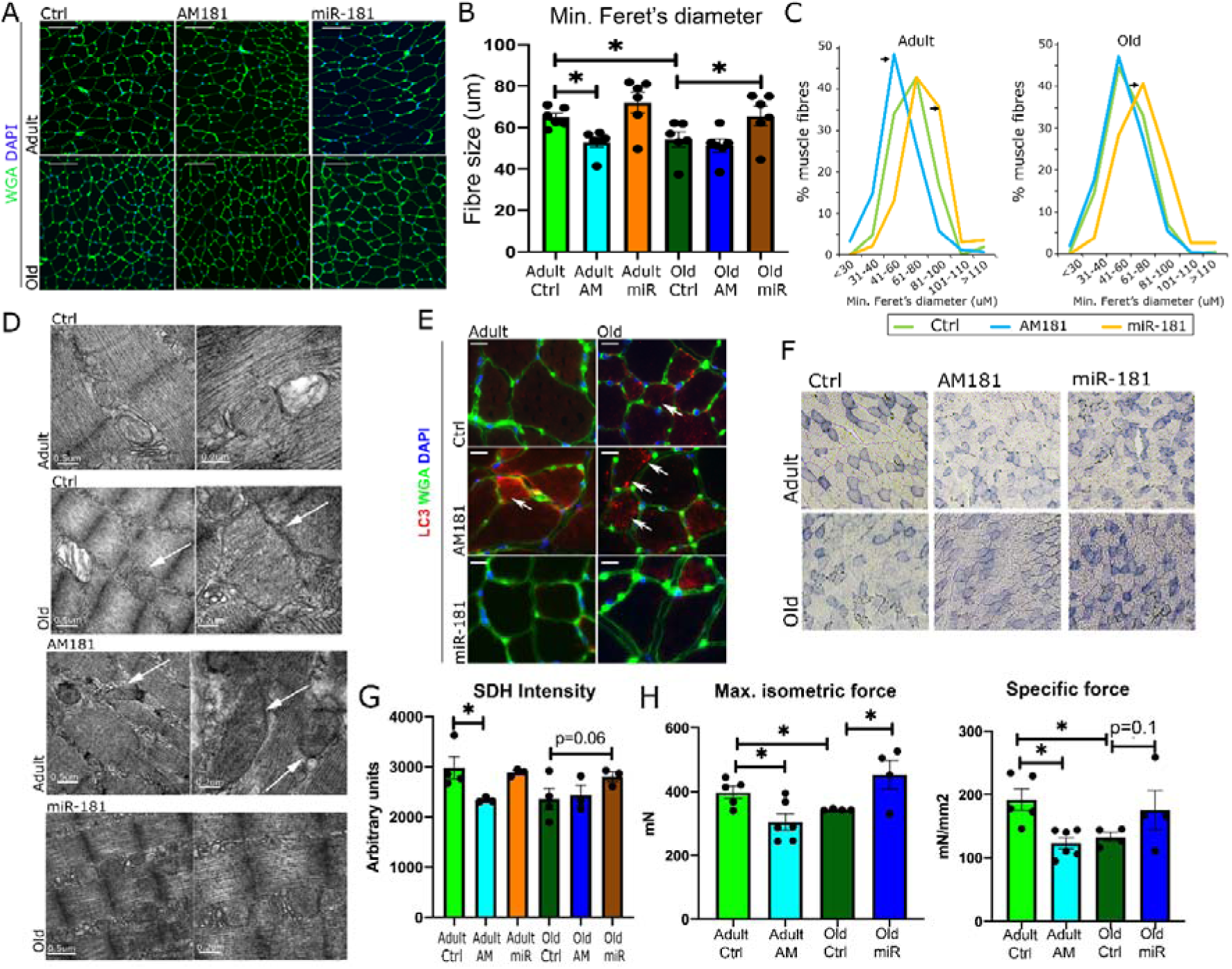
miR-181a regulates myofibre size and muscle strength *in vivo*. **A-C.** The effects of miR-181a on myofibre size in adult and old mice. WGA immunostaining (A), quantification of minimal Ferret’s diameter (B) and fibre distribution analysis (C) of TA of adult and old mice following intravenous injection of miR-181a mimic, antagomiR or saline (control). Scale bars indicate 100 µm; arrows indicate statistically significant changes in fibre distribution (Chi-square test, p<0.05). n=6; * - p<0.05 unpaired Student T test. **D.** Representative EM images of TA from adult and old mice. Arrows indicate abnormal mitochondria. Scale bars: 0.5 and 0.2 µm. **E.** Inhibition of miR-181a expression *in vivo* leads to the presence of more of LC3-positive myofibres (red), n=3, arrows indicate positive LC3 fibres. **F., G.** Histological examination of miR-181 effects on mitochondria: SDH staining and quantification (F) of TA of adult and old mice following miR-181 mimic, antagomiR-181 or saline injections, respectively. Scale bars indicate 200 µm; n=3. **H.** Muscle maximum and specific force following miR-181a gain- and loss-of-function *in vivo* evaluated in EDL muscles from adult and old mice; error bars indicate SEM; n = 4-6.

### miR-181a regulates mitochondrial quality, myofibre size and muscle force *in vivo*

We next investigated the physiological effects of miR-181a on muscle during ageing. Changes in miR-181a expression in adult and old mice had no significant effect on body or muscle weight (Fig. S6). However, miR-181a inhibition led to a decrease, whereas miR-181a overexpression led to an increase in myofibre size from adult and old mice respectively (Fig. 6A, B). The relative proportion of fibre sizes within the population of fibres revealed that miR-181a-treated mice had a higher proportion of larger fibres, whereas AM181a treatment resulted in a higher population of smaller fibres in muscle from adult mice (Fig. 6C). Moreover, a higher proportion of LC3-positive myofibres was detected in muscle of control old mice and AM181a-treated adult and old mice, with almost no LC3-positive fibres detected in adult saline and miR-181a-treated mice (Fig 6E). These results indicate miR-181a-mediated regulation of mitochondrial content is associated with fibre size.

The relative oxidative potential of muscle as an indication of mitochondrial function was decreased by miR-181a inhibition in the TA of adult mice, demonstrated by reduced intensity of succinate dehydrogenase (SDH) staining (Fig. 6F, G). miR-181a treatment of old mice resulted in a non-significant trend for increased SDH intensity (Fig. 6G). EM analysis of mitochondrial populations revealed the presence of swollen and mitochondria with disordered cristae, with some mitochondria expanding longitudinally between myofibrils, in muscle of control old and AM181a-treated adult mice. miR-181a treatment of old mice led to a lower proportion of structurally abnormal mitochondria and decreased the abundance of abnormally large mitochondria (Fig. 6D).

The physiological relevance of manipulating miR-181a expression was investigated by examining the effects of miR-181a on muscle force. Maximum and specific force of the EDL muscle was decreased during ageing (Fig. 6H) consistent with previous reports (Brooks and Faulkner, 1988). As miR-181a expression is decreased in muscle during ageing, we treated adult mice with AM181a to reduce miR-181a expression. Conversely, old animals were treated with miR-181a mimic in an effort to restore miR-181a expression. Maximum and specific force was decreased in muscle of adult mice treated with AM181a, whereas miR-181a overexpression in muscle of old mice restored maximum muscle force (Fig. 6H). Together, these data suggest that a decrease in miR-181a expression results in chronic dysregulated mitophagy in muscle during ageing associated with presence of abnormal mitochondria, loss of mitochondrial content and decline of muscle function.

In summary, a decrease in miR-181a expression resulted in increased expression of autophagy-associated genes, a decrease in fibre diameter and specific muscle force in adult mice mimicking ageing. Conversely, upregulating miR-181a levels in old mice improved mitochondrial morphology, increased fibre diameter and specific force. This data demonstrates microRNA-mediated fine-tuning of one of the key mechanisms associated with ageing, mitochondrial turnover, and provides proof-of-principle for the potential use of microRNA-based therapeutic approaches for ageing-associated disorders such as age-related muscle atrophy.

## Discussion

In the present study, we have demonstrated decreased mitochondrial content and quality associated with a dysregulated expression of autophagy and mitochondrial dynamics-associated proteins in skeletal muscle during ageing. Using bioinformatics and sensor constructs, we have identified miR-181a as a key regulator of autophagy-associated genes, controlling the expression of p62 and Park2, as well as indirect regulation of Park7/Protein DJ-1 expression and localisation. Changes in miR-181a levels and the concomitant changes in the expression of its direct and indirect targets result in altered myofibre size, mitochondrial content and quality and specific muscle force, suggesting miR-181a may be one of the key regulatory mechanisms underlying age-related muscle atrophy. This is the first report, to our knowledge, demonstrating a physiologically-relevant mechanism of miR-181a-mediated fine-tuning of mitochondrial dynamics on multiple levels through concerted regulation of expression of several autophagy-associated genes during ageing.

Mitochondrial quality has an important impact on the health and bioenergetics of healthy skeletal muscle and is tightly regulated by mitochondrial biogenesis, fusion, fission and selective degradation of mitochondria. Mitochondrial content decreases with age in skeletal muscle, however the exact mechanisms are not fully understood (Hepple, 2014). Functional autophagy and mitophagy are affected by ageing and have been associated with changes in mitochondrial dynamics (Carnio et al., 2014; Sebastian et al., 2016). We have detected an increase in the expression of the regulators of Pink1/Parkin dependent mitophagy during ageing and the presence of abnormal mitochondria, suggesting this mitophagy pathway is disrupted or ineffective, resulting in an accumulation of abnormal mitochondria.

The role of mitophagy and autophagy in skeletal muscle is context-dependent and even small changes in mitochondrial dynamics can have detrimental consequences on muscle mass and function with an accumulation of abnormal mitochondria (Carnio et al., 2014; Drake et al., 2016). Dysregulated mitophagy is associated with decreased functional mitochondria in muscle during ageing, where changes in mitochondrial fusion or fission may lead to the presence of abnormally large mitochondria which cannot be degraded via mitophagy (Mao and Klionsky, 2013). In *C. elegans* the age-related decline in mitophagy inhibits both the removal of dysfunctional mitochondria and impairs mitochondrial biogenesis (Palikaras et al., 2015). As miRs have been shown to fine-tune gene expression, they are ideal candidates to control this delicate between removal of dysfunctional/damaged mitochondria and mitochondrial biogenesis. Several microRNAs including miR-149, miR-761 and miR-494, have been identified that can inhibit or promote mitochondrial biogenesis ((Mohamed et al., 2014; Xu et al., 2015; Yamamoto et al., 2012) for review (Dahlmans et al., 2016).

In this study, we identified miR-181a, which has decreased expression in muscle during ageing, as a putative regulator of multiple genes associated with the regulation of mitochondrial dynamics. miR-181a mediated regulation of components of autophagy has previously been reported in ageing and cancer cell lines (Cheng et al., 2016; Rippo et al., 2014; Soriano-Arroquia et al., 2016a; Tekirdag et al., 2013) and recently overexpression of miR-181a/b-1 in chondrocytes resulted in improvements in mitochondrial metabolism (Zheng et al., 2019). Our results indicate miR-181a regulates autophagy/mitophagy and effector proteins, providing a mechanism of fine-tuning mitochondrial dynamics through parallel regulatory pathways. We have demonstrated that miR-181a can directly or indirectly regulate the expression and localisation/function of multiple autophagy-related genes: Park2, p62 and Protein DJ-1/Park7 (Figs. 3, 4 & 5) and concomitant with upregulation of mitochondrial biogenesis. Moreover, restored expression of miR-181a in the muscle of old mice prevented accumulation of mitophagy-associated proteins and was associated with increased mitochondrial content and improved mitochondrial quality (Figs. 5, 6). We also observed a decrease in the presence of swollen and abnormally large mitochondria in muscle of old mice following miR-181a overexpression (Fig. 6). Although these changes could be associated with miR-181a predicted target genes: Mfn1 and Mfn2, we did not observe consistent regulation of these genes by miR-181a at the mRNA level, suggesting an indirect regulation by miR-181a (Figs. S1, S5). Finally, we have shown that miR-181a regulates myofibre size and muscle force: adult and old mice treated with AM181a or miR-181a had significant changes in the expression of genes associated with mitochondrial dynamics, mitochondrial quality and muscle fibre diameter and specific force of muscle (Fig. 6).

In summary, we propose that miR-181a regulates skeletal muscle homeostasis and muscle metabolism, size and force, by regulating mitochondrial dynamics. miR-181a is downregulated during ageing and we propose that concomitant upregulation and accumulation of its mitophagy-associated targets paradoxically leads to a stalling or inhibition of mitophagy, resulting in the accumulation of damaged mitochondria. We have demonstrated that fine-tuning the levels of autophagy regulatory (p62) and mitophagy-associated proteins (DJ-1, Park2) and concomitant changes in the expression of Tfam, by overexpressing miR-181a, leads to improved mitochondrial dynamics resulting in increased myofibre size and muscle function. In this study we focused on the predicted and confirmed targets of miR-181a many of which are associated with the Pink/Parkin mitophagy pathway. It was therefore surprising to find that inhibition of Parkin, p62 and Protein DJ-1 by miR-181a overexpression resulted in increased mitophagy and mitochondrial biogenesis. Low levels of miR-181a in ageing muscle were associated with accumulation of its target proteins and stalled autophagy. Our data suggests that miR-181a regulates mitochondrial dynamics by fine-tuning the Pink/Park pathway and mitochondrial biogenesis, potentially in response to stress factors. Moreover, mitophagy is coordinated by a number of conserved cellular pathways, disruption of this balance can result in the accumulation of dysfunctional mitochondria and ageing (Palikaras et al., 2018). Exercise increases mitochondrial biogenesis in skeletal muscle, the increased expression of miR-181a in response to acute exercise (this study and (Safdar et al., 2009) suggests that miR-181a plays a key role in improving overall mitochondrial health. Our results indicate overexpression of miR-181a increases the expression of genes required for mitochondrial biogenesis and promotes mitophagy either directly or indirectly to restore mitochondrial dynamics.

Overall, the results of this study demonstrate for the first time the microRNA-mediated regulation of mitochondrial content in muscle during ageing with physiologically-relevant consequences on myofibre size and strength indicating the potential of microRNA-based therapies for age-related muscle atrophy.

## Materials and Methods

### Experimental Model and Subject Details

#### Mice

The study was performed using male wild type C57Bl/6 mice (adult: 6 months old; old – 22-24months old at the beginning of the treatment). Mice were obtained from Charles River (Margate). All mice were maintained under specific-pathogen free conditions and fed *ad libitum* a standard chow and maintained under barrier on a 12-h light-dark cycle. For miR-181a-5p expression manipulation, mice were injected with 2mg/kg body weight three times during 4-week period with miR-181a mimic (GE Healthcare, C-310435-05 conjugated to cholesterol) or custom antagomiR-181a (5’-FITC-mA*mC*mUmCmAmCmCmGmAmCmAmGmCmGmUmUmGmAmA*mT*mG*mU*mU - 3’Cholesterol; m – hydroxymethyl modified bases; * - phosphothioate bonds; CH – cholesterol) or saline (Control mice). *Extensor digitorum longus* (EDL) muscle force measurements and tissue collection were performed four weeks from the first injection. For tissue collection, mice were culled by cervical dislocation. The tissues were immediately excised, portions of TA muscles were either frozen and stored at −80°C or embedded in OCT, frozen in isopentane and stored at −80°C or for proteomics and western blotting homogenised. Experiments were performed in accordance with UK Home Office guidelines under the UK Animals (Scientific Procedures) Act 1986 and received ethical approval from the University of Liverpool Animal Welfare and Ethical Review Body (AWERB). For each experiment, n=3 (western blot and qPCR analyses) or n=6 (adult mice) n=4 (old mice; force measurements) mice were used.

#### Cell culture

C2C12 myoblasts were maintained in DMEM media (Sigma Aldrich, Dorset UK) supplemented with 10% Fetal Bovine Serum (FBS) and 1% penicillin/streptomycin (Sigma Aldrich, Dorset UK) (Goljanek-Whysall et al., 2012). For cholesterol-linked miR-181a mimic, antagomiR-181a or antagomiR scrambled, 50% confluent C2C2 myoblasts were treated at 100nM final concentration for 48h previous to treatment with 10 µM CCCP or 100nM Bafilomycin A (both Sigma Aldrich, Dorset UK). 16h following CCCP treatment or 1 hour after Bafilomycin A treatment, RNA and protein were isolated from treated cells or C2C12 cells were fixed in ice cold methanol and immunostained as described above (muscle staining). For mitophagy flux analyses, mitoQc reporter was used as previously (Allen et al., 2013). In brief, mitoQC viral molecules produced in HEK293T cells were used to transduce proliferating C2C12 cells. C2C12 cells were simultaneously treated with miR-181a mimic or antagomiRs and treated with DMSO (control), CCCP or Bafilomycin A and imaged as described above. Images were obtained using Zeiss fluorescent upright microscope (Model Imager M1) was used with AxioCam MRm and AxioVision Software version 4.8 and Olympus Fluoview3000 Laser Confocal microscope and analysed using Image J as described previously (Soriano-Arroquia et al., 2016b). Cytotoxicity and ATP generation were measured using mitochondrial ToxGlo assay (G8000) from Promega as per the manufacturer’s instructions.

### Method Details

#### *In situ* muscle function analysis

Force measurements of the EDL muscles were performed as described before via peroneal nerve stimulation in adult and old male mice (Sakellariou et al., 2016). Briefly, mice were anaesthetised using ketaset and dormitor. The distal tendon was secured to the lever arm of a servomotor (Aurora Scientific). Next, the knee was placed in a fixed position and the peroneal nerve exposed. Bipolar platinum wire electrodes were next placed on both sides of the peroneal nerve. Muscle optimal length (L_o_) was determined with during a series of 1□Hz stimulation and set at the length that generated the maximal force. EDL muscles were electrically stimulated to contract at L_o_ and optimal stimulation voltage (10□V) at 2□min intervals for 300□ms with 0.2□ms pulse width to determine maximum isometric tetanic force (P_o)_,. The frequency of stimulation was increased from 10 to a maximum of 300□Hz in 50□Hz increments. P_o_ was identified when the maximum force reached a plateau. Muscle fibre length (L_f_) and weight of EDL muscles were measured *ex vivo* and muscle cross-sectional area (CSA) was calculated (Sakellariou et al., 2016). Specific P_o_ (mN/mm^2^) was calculated by dividing P_o_ by total fibre CSA as described previously.

#### Muscle staining

TA muscles were cryosectioned at −20□°C through the mid-belly. 10□μm sections were rinsed with Phosphate Buffered Saline (PBS) and fixed in ice-cold methanol for 5□min. Sections were incubated in blocking solution (1%BSA: Sigma Aldrich, 10% normal horse serum: ThermoFisher Scientific, UK 0.3M glycine: Sigma Aldrich, Dorset UK, in 0.1% PBS-Tween: Sigma Aldrich) for 1 h at room temperature and next incubated with primary antibodies (p62, CoxIV: Cell Signaling,3E11, Tomm20:Abcam, UK, ab56783 (anti-mouse) or ab186734 (anti-rabbit), Lc3b: Abcam, UK, ab10912, DJ-1: Abcam, UK, ab76241, Parkin: Abcam, UK, ab15954, Pink1: Abcam, UK, ab23707 (all 1:1000 dilution in blocking buffer) diluted in 5% horse serum, 0.01% Triton X in PBS for 2h at room temperature. 3 PBS washes were performed and sections were next incubated with secondary antibodies diluted in 5% horse serum, 0.01% Triton X in PBS (anti-rabbit-Alexa647, anti-mouse – Alexa 532 or 488, Invitrogen, Paisley UK). Fluorescein labelled wheat germ agglutinin (WGA, Vector Laboratories, 5□μg/ml) was used to identify extracellular matrix. Nuclei were identified using 4′,6-diamidino-2-phenylindole (DAPI, 1□μg/ml, Sigma Aldrich, Dorset UK). SDH staining was performed as previously described (Smith et al., 2018). Transverse sections from 3–4 muscles/treatment group were examined by blinded observers to count the minimal Feret’s diameter. Image J software was used to analyse individual muscle fibres as described previously (Sakellariou et al., 2016).

#### Proteomics and Pathway analysis

TA muscles were immediately dissected and a portion homogenised directly in 50 mM ammonium bicarbonate pH 8. Protein lysates were prepared by centrifugation at 15,000g for 10min at 4°C and protein concentrations were calculated using Bradford assay (BioRad, UK) with BSA as a standard. 100µg of protein extract was diluted to 160µl with 25mM ammonium bicarbonate and denatured by the addition of 10µl of 1% RapiGest (Waters, Manchester, UK) in 25 mM ammonium bicarbonate and incubated at 80°C for 10min with shaking. 10µl of a 100mM solution of Tris(2-carboxyethyl)phosphine hydrochloride (TCEP) was added to reduce reversibly oxidised Cys residues followed by incubation at 60°C for 10min. Cys were then alkylated by addition of N-ethylmaleimide and incubated at room temperature for 30min. An aliquot of the samples was used at this point to check procedure by SDS PAGE. Proteolytic digestion was performed by addition of Trypsin (Invitrogen, Paisley, UK) followed by overnight incubation at 37°C. Digestion was terminated and RapiGest removed by acidification (3µl of TFA and incubated at 37°C for 45min) and centrifugation (15,000g for 15min).

##### LC−MS/MS and Label-Free MS Quantification

Samples were analysed using an Ultimate 3000 RSLC nano system (Thermo Scientific, UK) coupled to a QExactive mass spectrometer (Thermo Scientific, UK). 2µl of sample was diluted in 18 µl buffer (97% H_2_O, 3% MeCN and 0.1 % formic acid v/v) and 5µl corresponding to 250 ng of protein was loaded onto the trapping column (PepMap 100, C18, 75µm x 20mm) using partial loop injection for 7min at a flow rate of 4µL/min with 0.1% (v/v) TFA. Sample was resolved on the analytical column (Easy-Spray C18 75µm x 400mm, 2µm column) using gradient of 97% A (0.1% formic acid) and 3% B (99.9% ACN and 0.1% formic acid) to 60% A and 40% B over 120min at a flow rate of 300nL/min. Data dependent acquisition consisted of a 70,000 resolution full-scan MS scan (AGC set to 10^6^ ions with a maximum fill time of 250ms) and the 10 most abundant peaks were selected for MS/MS using a 17,000 resolution scan (AGC set to 5 × 10^4^ ions with a maximum fill time of 250ms) with an ion selection window of 3*m/z* and normalized collision energy of 30. Repeated selection of peptides for MS/MS was avoided by a 30sec dynamic exclusion window.

Label-free relative quantification was performed using PEAKS7 software (Bioinformatics Solutions Inc., Waterloo, Canada) and were searched against the UniProt mouse sequence database using an in house Mascot server (Matrix Science, London, UK). Search parameters used were: peptide mass tolerance 10ppm; fragment mass tolerance 0.01 Da, 1+, 2+, 3+ ions; missed cleavages 1; instrument types ESI-TRAP. Variable modifications included in search were: oxidation of methionine and cysteine residues and N-ethylmaleimide on cysteines. The mass spectrometry proteomics data have been deposited to the ProteomeXchange Consortium via the PRIDE (Vizcaino et al., 2016) partner repository with the dataset identifier PXD01009. Label free quantification data was imported into Perseus (Tyanova et al., 2016) software for generation of heatmaps. Heatmaps represent proteins that significantly change (>2 fold change in abundance −10log P value >20 (equivalent to a p value < 0.01) and identified with a False Discovery Rate of < 1%. Supporting information and identified proteins and their relative quantification. Pathway analysis of label free quantitative proteomic data was performed using PathVisio (Kutmon et al., 2015) together with WikiPathways (Kutmon et al., 2016) to visualize and highlight altered pathways from detected proteins.

#### Real-Time PCR and western blotting

RNA isolation and quantitative real time PCR (qPCR) were performed using standard methods (Goljanek-Whysall et al., 2014). cDNA synthesis (mRNA) was performed using 500ng RNA and SuperScript II (Invitrogen, Paisley UK) and cDNA synthesis (microRNA) was performed using 100ng RNA and miRscript RT kit II (Qiagen) according to the manufacturer’s protocol. qPCR analysis was performed using miRScript SybrGreen Mastermix (Qiagen) or sso-Advanced SybrGreen Mastermix (Biorad, Hercules USA) in a 10 µl reaction. Expression relative to β2-microblobulin or Rnu-6 and/or Snord-61 (microRNA) was calculated using delta delta Ct method. P-value was calculated using unpaired Student’s t-test. The sequences of primers used are included in Supplementary Table 1. For western blotting homogenised protein lysates were diluted using Laemmli buffer and separated on 12 % SDS PAGE gels. Briefly, 20 µg of protein was loaded, proteins were transferred using a semi-dry blotter, after transfer membranes were stained with Ponceau S to ensure equivalent loading. Membranes were blocked in 3% milk in TBS-T, following washing in TBS-T, membranes were incubated with primary antibodies (as above) at a dilution of 1 in 1000 in blocking buffer. Goat anti-rabbit HRP secondary antibody (Cell Signalling) was diluted 1 in 3000 in TBS-T. Thermo super signal west dura was used for chemiluminescence detection using a Chemidoc (BioRad), images were acquired and analysed using Image Lab 5.0 software (BioRad) and normalised using Ponceau S stain.

#### *In vitro* miRNA target prediction and validation

miR-181 targets were predicted using TargetScan v.6.2 (http://www.targetscan.org/, release June 2012) and miRWalk (http://zmf.umm.uni-heidelberg.de/apps/zmf/mirwalk2/) using mouse and human searches and broadly conserved microRNA target sites settings. 3’UTR regions with wild type or mutated miR-181a target site, were synthesised using GeneArt service (Invitrogen, Paisley UK) and cloned into a GFP TOPO vector (Invitrogen, Paisley UK). Sequences cloned are listed in Supplementary Table 2. C2C12 myoblasts were cultured in 96-well plates and transfected using Lipofectamine 2000^TM^ (Invitrogen, Paisley UK) with WT or mutant sensor (200 ng), with 100nM miR scrambled or miR-181a mimic (100nM; GE Healthcare) (Soriano-Arroquia et al., 2016a). Each experiment was carried out using at least two independent plasmid preparations in triplicates. GFP flourescence was measured 48h following transfections using FLUOstar Optima microplate reader (BMG Labtech).

#### Data Availability

The mass spectrometry proteomics data have been deposited to the ProteomeXchange Consortium via the PRIDE (Vizcaino et al., 2016) partner repository with the dataset identifier PXD01009. Source data for western blotting, qPCR primer sequences, sequences of 3’UTR cloned into GFP reporter construct sequences and label free proteomic data have been deposited into Mendeley data set doi:10.17632/gksmyhy89r.1.

#### Quantification and Statistical Analysis

##### Statistical analysis of SDH staining, myofibre size and muscle force

All data are represented as mean +/−_SEM. Data normality was assessed and unpaired Student’s t-test or Chi-square test, as indicated in figures, was performed using Prism version 5.01 software package for Windows (GraphPad Software, La Jolla California USA, www.graphpad.com). p-value < 0.05 was considered as statistically significant, n = 3-6 as indicated in figure legends.

##### qPCR Data

Expression relative to β2-microblobulin or Rnu-6 and/or Snord-61 (microRNA) was calculated using delta delta Ct method. p-value was calculated using unpaired Student’s t-test. W**estern blotting:** Band intensities were normalized using overall total protein intensity from Ponceau S staining. Relative expression was calculated using unpaired Student’s t-test with p-value < 0.05 considered significant. **Label free proteomics:** Proteins were quantified using top3 method using PEAKS7 label free software, proteins were considered significantly changes with p-value < 0.01 and > 2 fold change.

## Supplemental Information

Supplemental Information includes six figures.

## Acknowledgements

KGW and RMC were supported by Biotechnology and Biological Sciences Research Council grant (BB/L021668/1) awarded to KGW. The authors wish to thank the Electron Microscopy Laboratory, University of Liverpool and Marion Pope for EM analyses of muscle. The authors acknowledge the facilities and scientific and technical assistance of the Centre for Microscopy & Imaging at the National University of Ireland Galway (www.imageing.nuigalway.ie). Graphical abstract was produced using Affinity Designer version 1.6.4 (https://affinity.serif.com/en-gb/designer/).

## Author Contributions

Conceptualization, K.G.W. and B.M.D.; Methodology, K.G.W. and B.M.D.; Investigation, A.S.A., R.M.C., C.C., K.G.W. and B.M.D.; Writing – Original Draft K.G.W. and B.M.D.; Writing – Review & Editing, K.G.W., B.M.D., R.M.C. and A.S.A.; Funding Acquisition, K.G.W.; Supervision, K.G.W. and B.M.D.

## Contact for Reagent and Resource Sharing

Further information and requests for reagents may be directed to and will be fulfilled by the Lead Contacts, Brian McDonagh and Katarzyna Goljanek-Whysall (; or).

## Supplemental Materials

**Figure S1.** Expression of mitochondrial dynamics-associated genes in TA muscle of adult and old mice. Adult – 6 months old; old – 24 months old male C57BL6/J mice. Ex – TA following isometric contraction protocol. Expression relative to β2-microglobulin is shown. Representative images are shown. n=3. Error bars show SEM. * -p<0.05 Student’s t-test.

**Figure S2.** miR-181 does not regulate myoblast viability.

**A.** qPCR of miR-181a expression in C2C12 myoblasts following 10 µM CCCP treatment and transfections with scrambled antagomiR (control), miR-181a mimic or antagomiR-181a, respectively, relative to Rnu-6 expression.

**B.** C2C12 myoblasts treated with miR-181 mimic or anyagomiR-181 do not show decreased viability, however miR-181a treatment may improve mitochondrial function in the presence of CCCP, as compared to control. Mitochondrial ToxGlo assay (Promega) used.

**C. D.** Western blot and quantification of p62, DJ-1 and PARK-2 in C2C12 myoblasts treated with miR-181a mimic, antagomiR or control antagomiR (Scr) in control (DMSO) conditions or with CCCP treatment; representative western blots are shown.

*: p<0.05, unpaired Student T test. Error bars show SEM; n=3.

**Figure S3**. Immunostaining for TOM20 (mitochondrial marker) and mitophagy-associated proteins: LAMP1 and BNIP3 in C2C12 myoblasts following inhibition or overexpression of miR-181a in control (DMSO) or conditions promoting (CCCP) or inhibiting (BAF – bafilomycin) autophagy. Representative images are shown.

**Figure S4**. Negative control for immunostaining. Representative images of TAs negative controls (no primary antibody) immunostained for p62, DJ-1 and Park2 following miR-181a gain- and loss-of-function. Scale bars indicate 100 µm.

**Figure S5**. miR-181a gain- and loss-of-function in TA of adult and old mice leads to changes in the expression of several but not all mitochondrial dynamics-associated genes, relative to β2-microglobulin. Error bars show SEM * - p<0.05 Student T-test. Adult – 6 months old; old –24 months old male C57BL6/J mice; Scr - saline.

**Figure S6**. miR-181a mimic and antagomiR are effectively delivered into TA muscle via intravenous injections, however do not affect body weight or muscle mass. Changes in miR-181a expression in TA muscle of adult and old mice following intravenous injections of miR-181a mimic or antagomiR181a as compared to saline injected mice were detected by qPCR. Expression relative to Rnu-6. Error bars show SEM * - p<0.05 Student T-test. Adult – 6 months old; old –24 months old male C57BL6/J mice; Scr - saline.

