## Supplementary Files for "miR-181a Regulates p62/SQSTM1, Parkin and Protein DJ-1 Promoting Mitochondrial Dynamics in Skeletal Muscle Ageing"

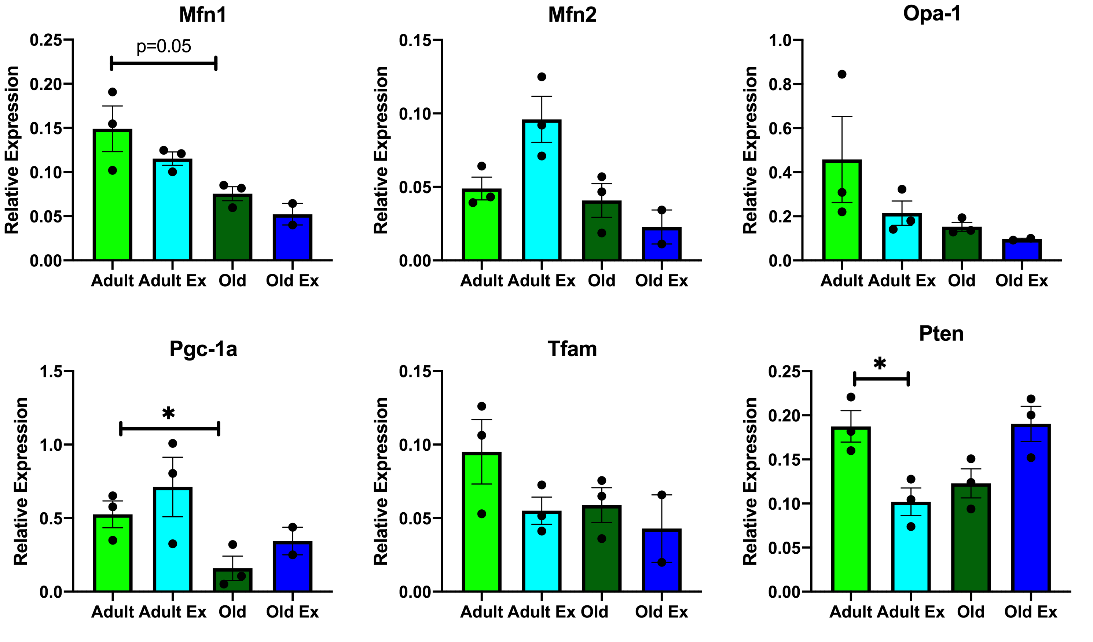

**Figure S1.** Expression of mitochondrial dynamics-associated genes in TA muscle of adult and old mice. Adult – 6 months old; old – 24 months old male C57BL6/J mice. Ex – TA following isometric contraction protocol. Expression relative to β2-microglobulin is shown. Representative images are shown. n=3. Error bars show SEM. * - p<0.05 Student’s t-test.

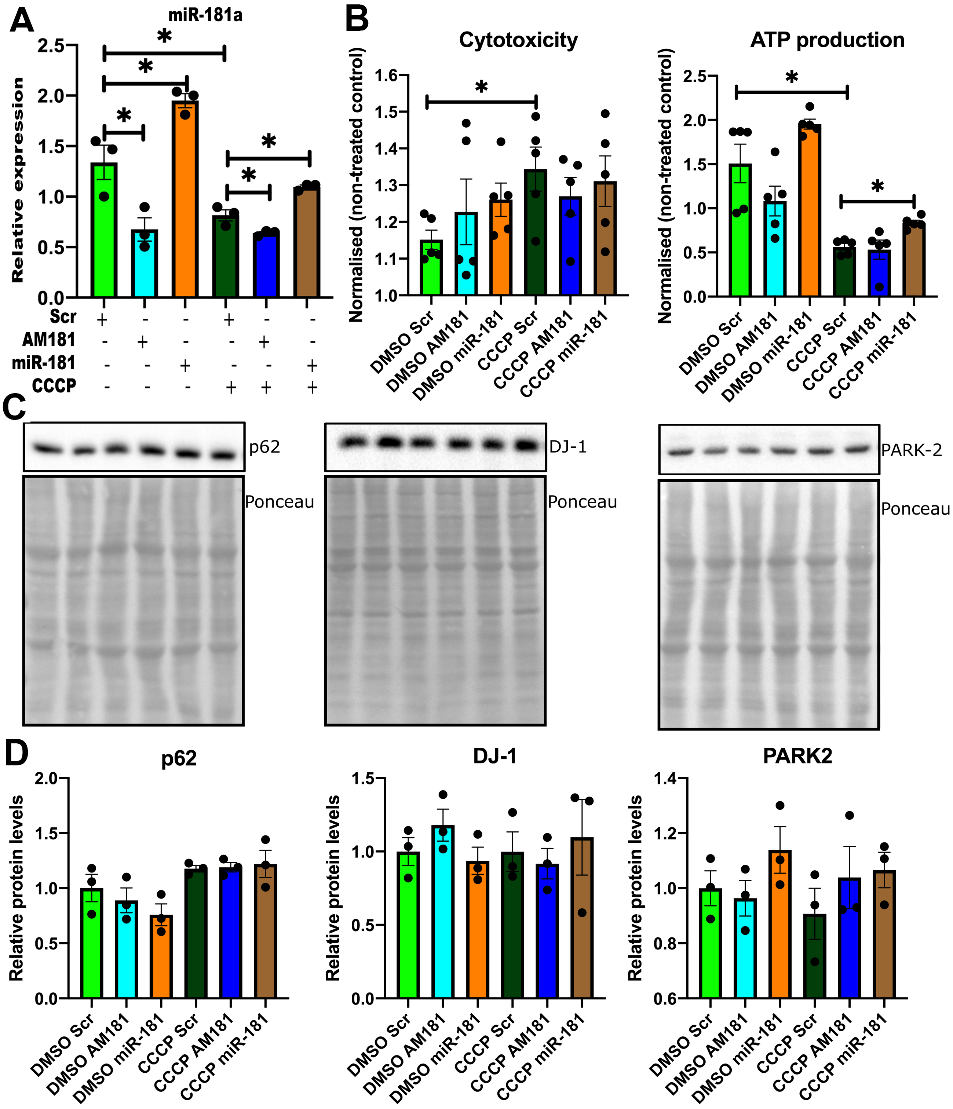

**Figure S2. miR-181 does not regulate myoblast viability.**

**A.** qPCR of miR-181a expression in C2C12 myoblasts following 10 µM CCCP treatment and transfections with scrambled antagomiR (control), miR-181a mimic or antagomiR-181a, respectively, relative to Rnu-6 expression.

**B**. C2C12 myoblasts treated with miR-181 mimic or anyagomiR-181 do not show decreased viability, however miR-181a treatment may improve mitochondrial function in the presence of CCCP, as compared to control. Mitochondrial ToxGlo assay (Promega) used.

**C. D.** Western blot and quantification of p62, DJ-1 and PARK-2 in C2C12 myoblasts treated with miR-181a mimic, antagomiR or control antagomiR (Scr) in control (DMSO) conditions or with CCCP treatment; representative western blots are shown.

*: p<0.05, unpaired Student T test. Error bars show SEM; n=3.

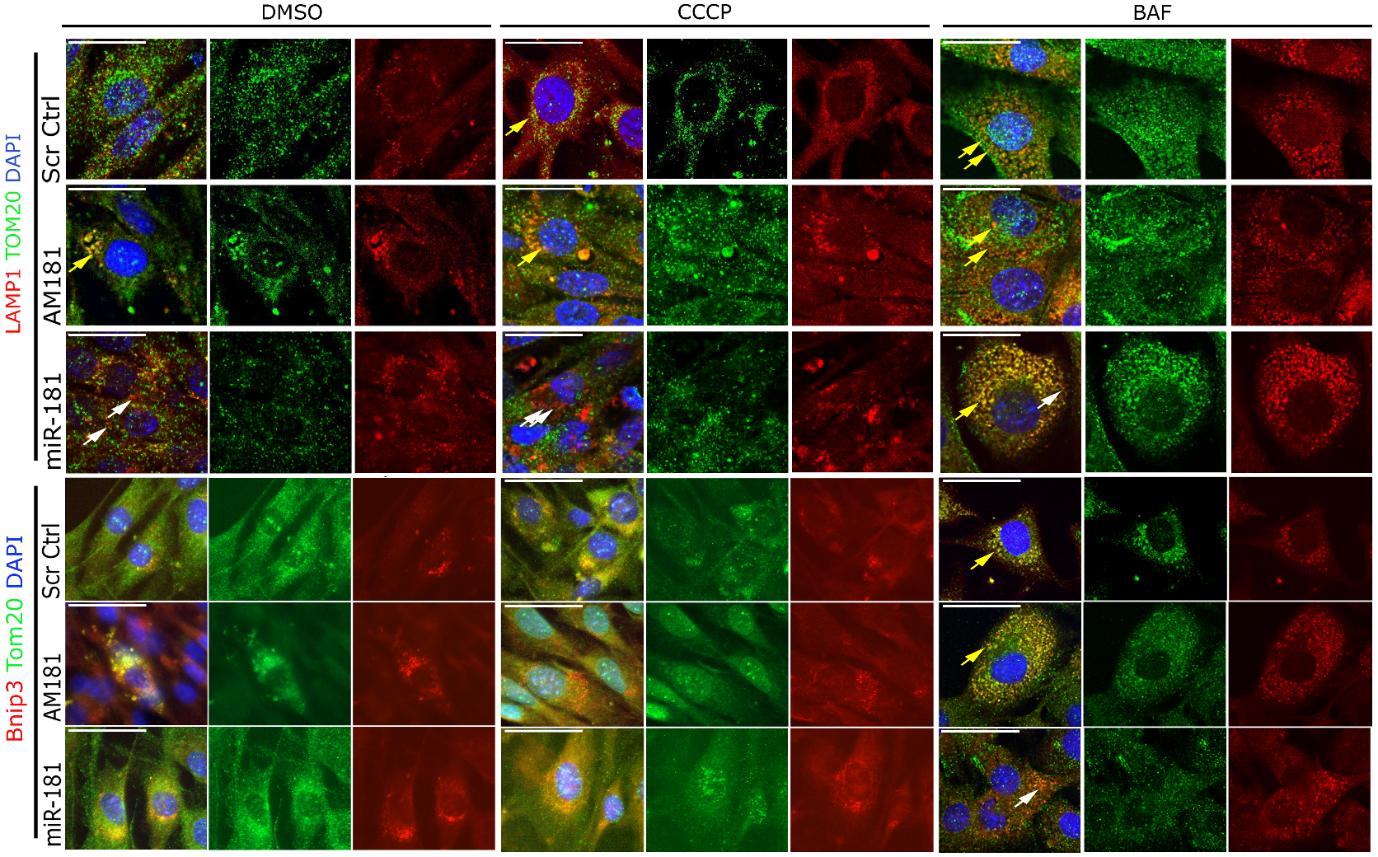

**Figure S3**. Immunostaining for TOM20 (mitochondrial marker) and mitophagy-associated proteins: LAMP1 and BNIP3 in C2C12 myoblasts following inhibition or overexpression of miR-181a in control (DMSO) or conditions promoting (CCCP) or inhibiting (BAF – bafilomycin) autophagy. Representative images are shown.

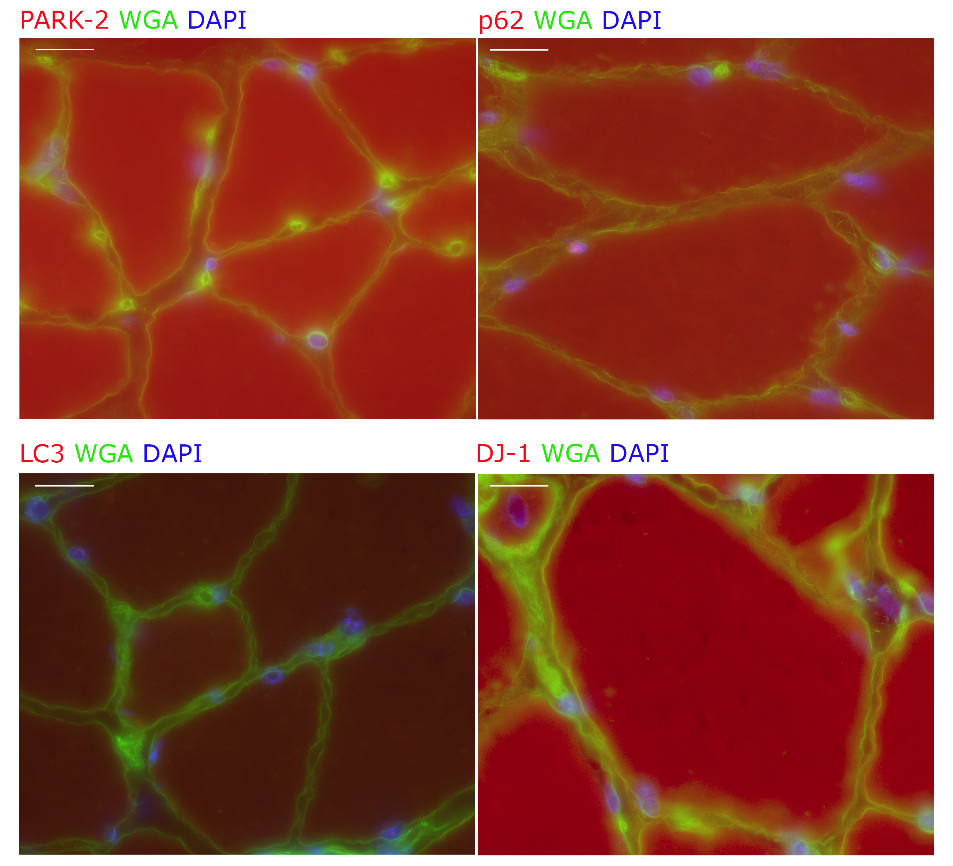

**Figure S4**. Negative control for immunostaining. Representative images of TAs negative controls (no primary antibody) immunostained for p62, DJ-1 and Park2 following miR-181a gain- and loss-of-function. Scale bars indicate 100 µm.

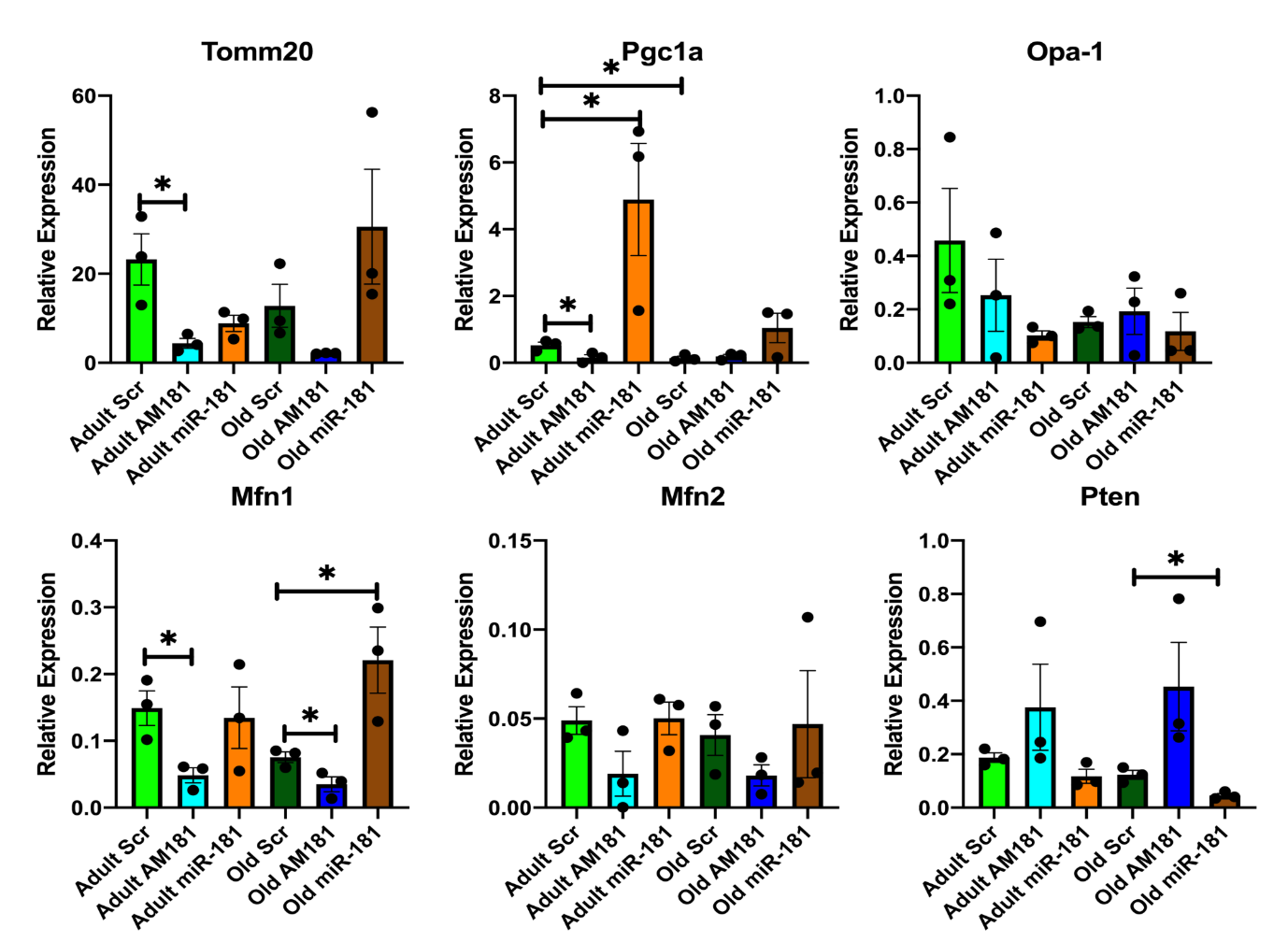

**Figure S5**. miR-181a gain- and loss-of-function in TA of adult and old mice leads to changes in the expression of several but not all mitochondrial dynamics-associated genes, relative to β2-microglobulin. Error bars show SEM * - p<0.05 Student T-test. Adult – 6 months old; old –24 months old male C57BL6/J mice; Scr - saline.

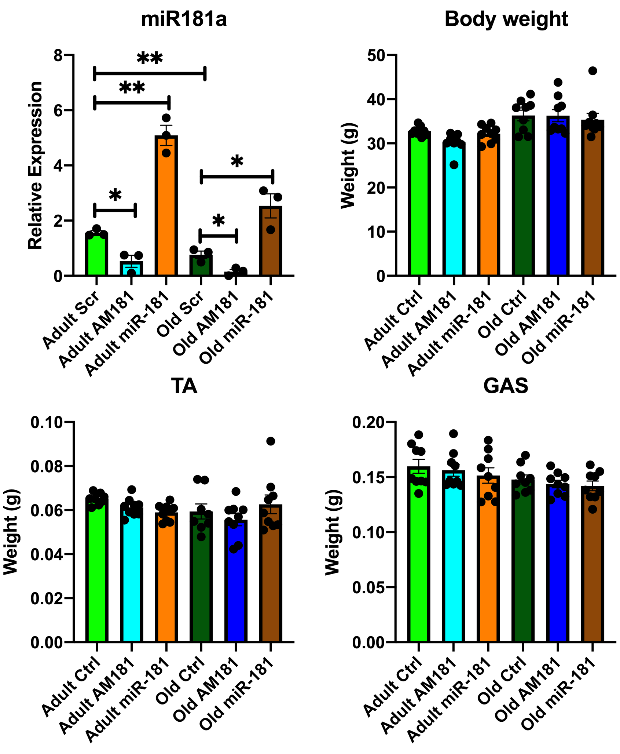

**Figure S6**. miR-181a mimic and antagomiR are effectively delivered into TA muscle via intravenous injections, however do not affect body weight or muscle mass. Changes in miR-181a expression in TA muscle of adult and old mice following intravenous injections of miR-181a mimic or antagomiR181a as compared to saline injected mice were detected by qPCR. Expression relative to Rnu-6. Error bars show SEM * - p<0.05 Student T-test. Adult – 6 months old; old –24 months old male C57BL6/J mice; Scr - saline.
